# The Creation of A Negative Slope Conductance Region by the activation of The Persistent Sodium Current Prolongs Near-Threshold Synaptic Potentials

**DOI:** 10.1101/084723

**Authors:** Cesar C. Ceballos, Antonio C. Roque, Ricardo M. Leão

## Abstract

A change of the input resistance (R_in_) of the neuron involves a change in the membrane conductances by opening and closing of ion channels. In passive membranes, i.e., membranes with only linear leak conductances, the increase or decrease of these conductances leads to a decrease or increase of the R_in_ and the membrane time constant (τ_m_). However, the presence of subthreshold voltage dependent currents can produce non-linear effects generating deviations from this relationship, especially the contradictory effect of negative conductances, as produced by the sodium-persistent current (I_NaP_), on the R_in_. In this work we aimed to analyze experimentally and theoretically the impact of the negative conductance produced by I_NaP_ on R_in_. Experiments of whole-cell patch-clamp conducted in CA1 hippocampus pyramidal cells from brain slices showed a paradoxical voltage-dependent decrease of the R_in_ and the τ_m_ in subthreshold membrane potentials close to the firing threshold after the perfusion with TTX, which inhibits I_NaP_. This effect is postulated to be a result of the negative slope conductance in the subthreshold region produced by this conductance. The analysis of the experimental data, together with simulations found that the slope conductance of I_NaP_ is negative for subthreshold membrane potentials and its magnitude is voltage dependent in the same range observed for the voltage-dependence of R_in_ and τ_m_. The injection of an artificial I_NaP_ using dynamic-clamp in the presence of TTX restored the R_in_ and τ_m_ to its original values. Additionally the injection of an artificial leak current with a negative conductance in the presence of TTX restored the R_in_ and τ_m_ as the artificial I_nap_ did. On the other hand, the injection of an artificial leak current with a positive conductance in the presence of TTX had no effect on the R_in_ and τ_m_. We conclude that I_NaP_ increases the R_in_ and τ_m_ by the negative slope conductance observed in its non-monotonic I-V relationship. These results demonstrate that the effect of I_nap_ on R_in_ and τ_m_ is stronger in potentials near the firing threshold, which could potentiate the temporal summation of the EPSPs increasing their temporal integration and facilitating action potential firing. Because of its negative slope conductance, I_NaP_ is more effective in increasing excitability near threshold than a depolarizing leak current.

## INTRODUCTION

The membrane input resistance (R_in_) and membrane time constant (τ_m_) are traditionally view, based on the cable theory, as a result of the passive non-voltage dependent leak or background channels (Enyedi et al., 2010, Dallérac et al., 2014, Du et al., 2016), and not depending on the voltage-dependent channels (Koch 1998). However, the electrical behavior of neurons is never purely passive because most neurons also express voltage dependent currents activated at subthreshold membrane potentials making R_in_ and τ_m_ voltage-dependent (Surges et al., 2004, Nisenbaum et al., 1995, Klink et al., 1997, Fernandez et al., 2015, Yamada-Hanff & Bean, 2015). Thus, the classical cable theory is insufficient to explain the subthreshold membrane properties of the soma.

The persistent sodium current (I_NaP_) is a subthreshold non-inactivating voltage dependent current which is expressed in several neuronal types (Stafstrom et al., 1985; Crill, 1996; French et al., 1999; Magistretti et al., 1999; Del Negro et al., 2002; Leão et al., 2005, 2012). I_NaP_ is rapidly activated and deactivated (τ < 250 μs, Carter et al., 2012), but has a slow inactivation, remaining active for hundreds of milliseconds (French et al., 1990). I_NaP_ is not generated by a different type of Na_V_ subunit, but depends on the interaction of the N_aV_ alpha subunit with beta subunits (Qu et al., 2001; Aman et al., 2009). I_NaP_ affects substantially the neuronal excitability boosting depolarization near the threshold and producing spontaneous firing (Leao et al., 2012, Raman et al., 2000, Pennartz et al., 1997) and mutations in N_aV_ subunits increasing the expression of I_NaP_ are related to pathologies as epilepsies (reviewed by Stafstrom, 2007). The voltage-dependence of I_NaP_ is similar to the classical voltage dependent sodium current, with a non-monotonic relationship originated from the interaction between its channel activation kinetics and its reversal potential (Izhikevich 2007, Ghigliazza & Holmes, 2004). This implies that the current-voltage (IV) relationship of I_NaP_ displays both a positive and a negative slope conductance, which are determined by dI/dV at the different points of the curve, being the negative conductance expressed at the activation phase of the curve (Izhikevich 2007, Stafstrom et al., 1982, Yamada-Hanff & Bean 2015, Ghigliazza & Holmes, 2004). Negative slope conductances increase R_in_ and τ_m_, despite to the increased channel conductance (Fernandez et al., 2015) and experimental evidence shows that I_NaP_ increases R_in_ and τ_m_ in its negative slope conductance region (Wilson 2005, Haj-Dahmane et al., 1996, Yamada-Hanff et al., 2013, Fernandez et al., 2015, Economo et al., 2014, Yamada-Hanff et al., 2015, Jacobson et al., 2005, Yaron-Jakoubovitch et al., 2008, Boehlen 2012, Waters et al., 2006, Buchanan et al., 1992).

For more than three decades it is known that the amplitude and duration of excitatory post-synaptic potentials (EPSPs) are voltage dependent, and that depolarizing the membrane potential to values close the action potential threshold amplifies its amplitude and prolongs its decay phase (Deisz et al., 1991, Thomson et al., 1988, Stafstrom et al., 1985). In cortical and hippocampal pyramidal cells, these effects are mediated mainly by I_NaP_ (Stuart et al., 1995; Fricker et al., 2000). The mechanism for the amplification of the EPSP amplitude is attributed to the fast activation of the I_NaP_ that boosts depolarization and increases R_in_ by its negative slope conductance (Yamada-Hanff & Bean 2015). However no satisfactory explanation has been provided for the prolongation of the EPSP. From an intuitive point of view, the slow inactivation of I_NaP_ could explain the longer duration of EPSPs near threshold. However, the non-inactivating I_h_ current shortens the duration of EPSP (Magee 1988). Both, I_NaP_ and I_h_ are depolarizing and non-inactivating currents, suggesting that neither aspect is responsible for the EPSP prolongation. Furthermore, I_NaP_ has a fast deactivation whereas I_h_ has slow deactivation, strongly suggesting that channel deactivation kinetics is not involved in the prolongation of EPSPs. Recently, Yamada-Hanff & Bean (2015) have shown that I_NaP_ increases the τ_m_ whereas I_h_ decreases the τ_m_, and that the increase or decrease of τ_m_ matches with regions of negative and positive slope conductance of the IV curves, respectively. These observations suggest that the mechanism underlying the prolongation of near threshold EPSP is possibly related to the negative slope conductance produced by I_NaP_ in the subthreshold membrane potentials.

In this work we aimed to establish the biophysical mechanisms underlying the prolongation of the EPSP decay in CA1 pyramidal cells. For this we develop a new analytical solution for the steady-state slope conductance of non-inactivating voltage dependent currents in order to gain some insight on the mechanisms by which I_NaP_ activation increases R_in_ and τ_m_. Mathematical, computational and experimental approaches were used to establish the mechanism of the increasing R_in_ and τ_m_ by the negative slope conductance of the I_NaP_. To further investigate the relationship between R_in_, τ_m_ and negative slope conductance in real neurons, we applied an artificial I_NaP_ to hippocampal pyramidal cells using dynamic clamp. Finally, we investigate the effect of a pure negative linear conductance on the R_in_, τ_m_ and the prolongation of EPSPs. Our hypothesis was that the ability for the I_NaP_ to increase R_in_, τ_m_ and duration of EPSPs is related only to its negative slope conductance and not dependent on the channel kinetics or ionic conductance. Our experimental and theoretical results show that the negative conductance of I_NaP_ is enough to increase the τ_m_ and to prolong the decay phase of EPSPs.

### METHODS

#### Hippocampal slices and electrophysiology

The experimental procedures were approved by the Ethics Committee on Animal Experimentation (CETEA) of the Medical School of the University of Sao Paulo at Ribeirao Preto (FMRP-USP). Male Wistar rats (P18 to P22) were anesthetized with isoflurane and decapitated and their brains removed. Coronal slices (300 μm thickness) containing the dorsal hippocampus were obtained with a vibratome (Vibratome 1000 Plus) in an ice-cold solution containing (mM): 87 NaCl, 2.5 KCl, 1.25 NaH_2_PO_4_, 25 NaHCO_3_, 0.2 CaCl_2_, 7 MgCl_2_, 25 dextrose and 75 sucrose, pH 7.4 when oxygenated with 95% O_2_/5% CO_2_. The slices were incubated in an artificial cerebrospinal solution (aCSF) during 45 min at 35^o^C and thereafter at room temperature (25^o^C). The aCSF contained (mM): 120 NaCl, 2.5 KCl, 1.25 NaH_2_PO_4_, 25 NaHCO_3_, 2 CaCl_2_, 1 MgCl_2_ and 20 dextrose, pH 7.4 when oxygenated with 95% O2/5% CO2 (osmolality ~310 mOsmol/kg H_2_O). CA1 pyramidal cells (PYR) were visualized with a microscope (Olympus BX51W) equipped with DIC-IR optics using a 40x water immersion objective. Recordings were obtained with borosilicate microelectrodes presenting a resistance between 4-6 MΩ when filled with the internal solution (mM): 138 K-gluconate, 8 KCl, 10 Na-phosphocreatine, 10 HEPES, 0.5 EGTA, 4 Mg-ATP, 0.3 Na-GTP, pH 7.4 adjusted with KOH and 295 mOsmol/kg H_2_O.

Pyramidal cells were identified based on the location on the CA1 pyramidal layer, firing behavior and the triangular shape of the soma. Whole-cell patch-clamp recordings were conducted in a controlled temperature around 33-36 ^o^C using an in-line heater (Scientifica). Series resistance was monitored through the experiments from the size of the capacitive transient in response to a 5 mV voltage step in voltage clamp mode. Recordings with access resistance larger than 20 MΩ were discarded. Series resistance was compensated at 80%. All the recordings were done in the presence of the GABA_A_ antagonist picrotoxin (PTX, 20 μM). Tetrodotoxin (TTX, 100 nM) was perfused to block the persistent sodium currents, when indicated. Data were acquired using PATCHMASTER (HEKA) at a 40 KHz rate, filtered with a low pass filter (Bessel, 3 kHz) using an EPC 10 amplifier (HEKA). Drugs were prepared in aliquots 1000 x concentrated and diluted into the perfusion solution in the day of the experiment. PTX was obtained from Sigma and TTX was obtained from Alomone Labs.

#### Dynamic-clamp

Our dynamic-clamp was implemented in a HP notebook running Linux with RTAI (RealTime Application Interface for Linux). Comedi library packages allowed the control of a DAQ board (Texas Instrument) plugged to the EPC 10 amplifier. Codes were written in C and the equations were solved using first order exponential Euler method with 50 μs steps (Dagostin al, 2015).

Dynamic-clamp recordings were done recording the membrane potential and injecting current onto the cell in the whole-cell current clamp mode through the same pipette. An artificial I_NaP_ was injected in neurons bathed in standard aCSF perfusion and in aCSF with TTX. Artificial I_NaP_ was calculated and injected in real time using the equations:

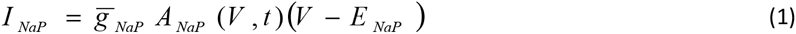

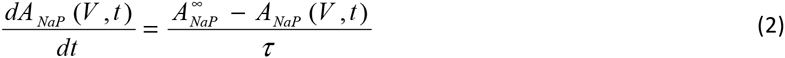

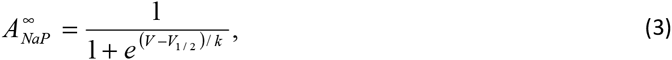

where the maximum conductance 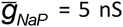, the reversal potential *E*_Na_ = +50 mV and the activation time constant *τ* = 2 ms. The parameters of the steady state activation variable 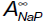 were fitted to reproduce the persistent sodium channel kinetics (V_1/2_ = -50 mV, k = 6) in agreement with experimental data from Yamada-Hanff et al., (2015). Membrane potential and currents were sampled at a 20 KHz rate. Membrane potential was recorded and corrected on-line for the liquid junction potential calculated in 10 mV after the equations were solved.

Artificial linear conductances were simulated using dynamic clamp. We tested two linear conductances: one with a positive conductance and the other with a negative conductance (Bose et al., 2014; Fernandez et al., 2015), following the equation 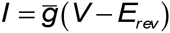 where 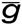 is the maximum conductance in nS and *E*_rev_ is the reversal potential in mV. For the current with positive conductance we used 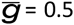 and *E*_rev_ = 50, and for the current with negative conductance we used 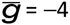 and *E*_rev_ = -80. These conductance values were chosen from the NaP slope conductance and chord conductance curves for the membrane potential −70 mV (see Figure 1B and Figure 1C). In the experiments when the negative conductance was injected, we restricted the current injection to the range between -80 mV and -60 mV in order to prevent instabilities.

#### Data analysis

Measured liquid junction potential was of 10 mV and was corrected off-line. Experiments in the voltage-clamp mode were conducted at a holding potential of -80 mV. In order to isolate the I_NaP_, voltage ramps were applied from -90 mV to -30 mV for 4 seconds. I_NaP_ was obtained by applying TTX (100 nM) and subtracting the currents produced by the ramps before and after TTX.

Input resistance was measured in current-clamp mode analyzing voltage-current (VI) relationships, before and after application of TTX. VI curves were obtained injecting 1 s current pulses with +10 pA steps from -400 pA to +400 pA. The steady state potential was measured during the last 100 ms of the pulse. The input resistance for each membrane potential was determined from the slope of the linear regression using 3 consecutive steady state potentials from the VI curve. Membrane time constant was measured in the current-clamp mode by fitting a single exponential to the first 100 ms of the voltage change in response to a small (+20 pA) current injection. When measuring the input resistance and the membrane time constant, 7 small current pulses were injected in order to maintain ΔV as minimum as possible. In order to determine the effect of the I_NaP_ on the EPSP shape (amplitude and duration), artificial EPSC (aEPSC) were injected through the recording pipette while the membrane potential was changed. aEPSCs were built using two consecutive current ramps, (2 ms rising and 5 ms decay). The maximum current amplitude (200 pA) was set in order to obtain ~ 5 mV aEPSPs at −90 mV.

We measured the amplitude (mV) and the area under the aEPSP (mV*ms) in order to determine the amplification in the amplitude and the decay time of aEPSPs by I_NaP_. Area normalized by the amplitude was also determined to isolate the prolongation of the aEPSP decay. Due to the variability of the shape of the aEPSPs near the firing threshold, we were unable to fit the aEPSP decay using exponential functions, and for this reason we decided to measure the area of the aEPSP to estimate its decay.

Data were analyzed using algorithms written in Igor Pro (Wavemetrics) and Matlab. Data were presented as mean ± SEM. Means were statistically compared using One way repeated measures ANOVA, Two way repeated measures ANOVA and linear regression in GraphPad Prism (GraphPad Software, La Jolla, CA). Statistical significance was set below *p* = 0.05.

#### Neuron model

We considered a single compartment model neuron with only linear (*I*_L_) and non-inactivating voltage dependent currents (*I*_NI,j_, *j* = 1, …, *n*), where *n* is the number of different non-inactivating currents. The membrane voltage is described by the equation

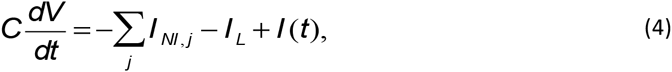

where *C* is the membrane capacitance and *I*(*t*) is the injected current. In the steady state this equation implies

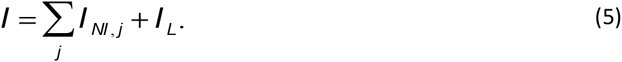

Using the Hodgkin-Huxley formalism to represent the ionic currents in the steady state one gets

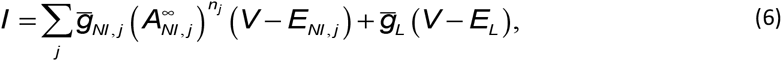

where 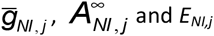 and *E_NI, j_* represent, respectively, the maximum conductance, the steady-state activation variable and the reversal potential of the *j*th non-inactivating current type. 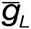 and *E_L_* are, respectively, the maximum conductance and the reversal potential of the linear current, and *n_j_* are natural numbers introduced to fit the experimental data. The activation variables are represented by Boltzmann terms like the one in equation (3). In this equation, V_1/2_ is the membrane potential at which 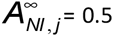 and k is the slope factor, which is a real number whose magnitude represents the slope of the sigmoid activation curve. Positive k means that the current is activated by hyperpolarization and negative k means that the current is activated by depolarization.

#### Slope and chord conductance/resistance

The infinitesimal or differential definition of the input conductance in the steady state *G*_in_ is

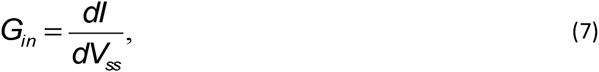

where *V*_ss_ is the steady-state membrane potential. This definition is commonly known as the slope conductance, since it corresponds to the slope of the steady-state *I-V* plot. The slope conductance at the steady state of a non-inactivating voltage-dependent current is obtained by differentiating its current equation

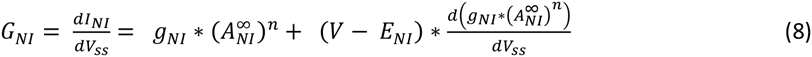

Splitting the equation (8) in two terms

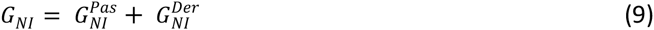

Where 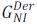 is defined as the derivative term of the slope conductance and 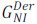 is defined as the passive term of the slope conductance. Notice that 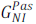 corresponds to the so called chord conductance or to the biophysical conductance of the channel, so that 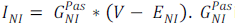 is always positive, whereas 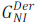 could be positive or negative.

The input conductance of a membrane that only contains ionic linear currents (I_L_) and ionic non-inactivating voltage dependent currents (I_NI_) is obtained differentiating the equation (6) with respect to V:

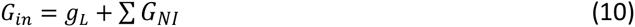

Thus, the input conductance in the steady state corresponds to the sum of the slope conductance of each ionic current in the steady state. Since the slope conductance characterizes the response of the membrane current to small changes in the membrane potential, its inverse is the slope resistance, or input resistance:

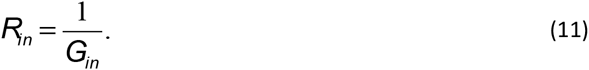

Both the input resistance or the slope conductance can be used to characterize membranes and/or single channels. However, these definitions, although operationally straightforward, do not meet the criteria for an Ohmic conductance. From the viewpoint of Ohm's law, a more appropriate definition of the channel conductance is the one in which the conductance at a point on the steady-state *i-V* trajectory is defined by the slope of a straight line connecting that point with the reversal potential. This conductance, typically referred to as the *chord* conductance, is defined on the basis of Ohm's law as

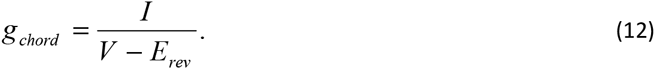

So, the chord conductance corresponds to the biophysical conductance of the channel. Slope conductance and chord conductance are the same for all voltage only when the I-V curve is a straight line, i.e, when the current is linear or passive, like produced by leak channels.

#### Membrane time constant

In a membrane containing only linear currents, the membrane equation can be written as

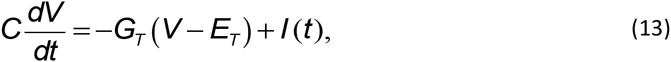

where 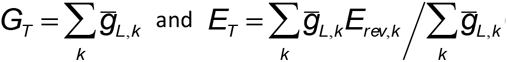 (*k* is the index for the different linear currents). The time evolution of *V*(*t*) for a membrane with linear currents only is described by the following equation (Koch 1999, Sterratt et al., 2011):

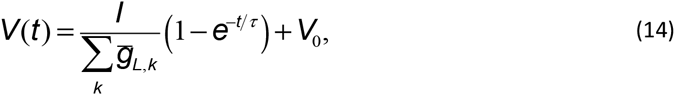

where *V*_0_ satisfies the equation 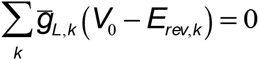. The membrane time constant τ in equation (14) is defined as

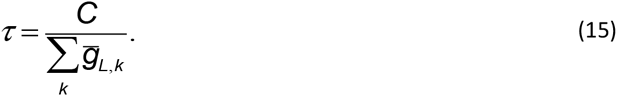

The single compartment was a cylinder with length and basis diameter 70 μm. The maximum conductance and reversal potential of the *I*_NaP_ current were, respectively, 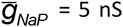 and *E*_Na_ = 50 mV. The maximum conductance and reversal potential of the *I*_L_ current were, respectively, 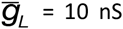 and *E*_L_ = −90 mV. Simulations were run both in NEURON and MATLAB.

## RESULTS

### The negative slope conductance of I_NaP_ predicts an increase in τ_m_ by increasing the R_in_

First we show analytically how an ohmic leak current and I_NaP_ determine R_in_ and τ_m_. I_NaP_ has a fast activation kinetics when compared with the membrane τ_m_, i.e., the channel time constant (τ_NaP_) is much faster than τ_m_, (τ_NaP_ << τ_m_). Since the I_NaP_ activation kinetics is almost instantaneous (Carter et al., 2012), its steady state slope conductance (G_NaP_) equals its instantaneous slope conductance. Thus, it is possible to rewrite the I_NaP_ current equation (1) as a linear current with a slope conductance equal to G_NaP_ (see Fig 1A).

So that;

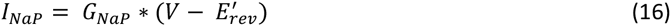

where the new reversal potential E^'^_rev_ is now voltage dependent, i.e., E^'^_ver_ = E^'^_ver_(V). However, if a small current ΔI is injected, sufficiently small to assume that G_NaP_(V) = G_NaP_(V_ini_) and E^'^_rev_(V) = E^'^_rev_(V_ini_), in other words, both G_NaP_ and E^'^_rev_ don’t change and are independent of the membrane potential and time, so;

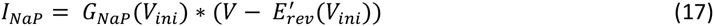

what now has the format of an ohmic leak current. Thus, I_NaP_ behaves as a linear current when ΔI is very small, and now its chord conductance is equivalent to G_NaP_. Thus, the time evolution of V = V(t) for a membrane with a linear leak current and I_NaP_ with fast activation follows equation 14, and τ_m_ is defined as in equation 15 (see Methods), where

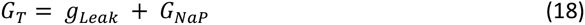

summarizing, the contribution of I_NaP_ to the τ_m_ is via its slope conductance G_NaP_. So that, the I_NaP_ negative slope conductance increases τ_m_ decreasing G_T_, meanwhile the linear currents decrease τ_m_ increasing G_T_.

Figure 1 shows computational simulations with a single compartment neuron containing a linear current and I_NaP_. Figure 1B and 1C show that both voltage change and voltage rise rate are dependent to membrane potential, being bigger and slower as membrane potential depolarizes. Figure 1D and 1E show that R_in_ and τ_m_ are voltage dependent and increase with depolarization. When we correlate R_in_ with τ_m_ for all the membrane potentials, it is clear that τ_m_ is directly proportional to R_in_ (Figure 1F). This result predicts that the mathematical relation τ_m_ = C* R_in_ is valid for both a passive membrane (only expressing leak channels) and a membrane containing leak and NaP channels. In the next sections we validate our predictions through experimental observations.

**Fig 1.**
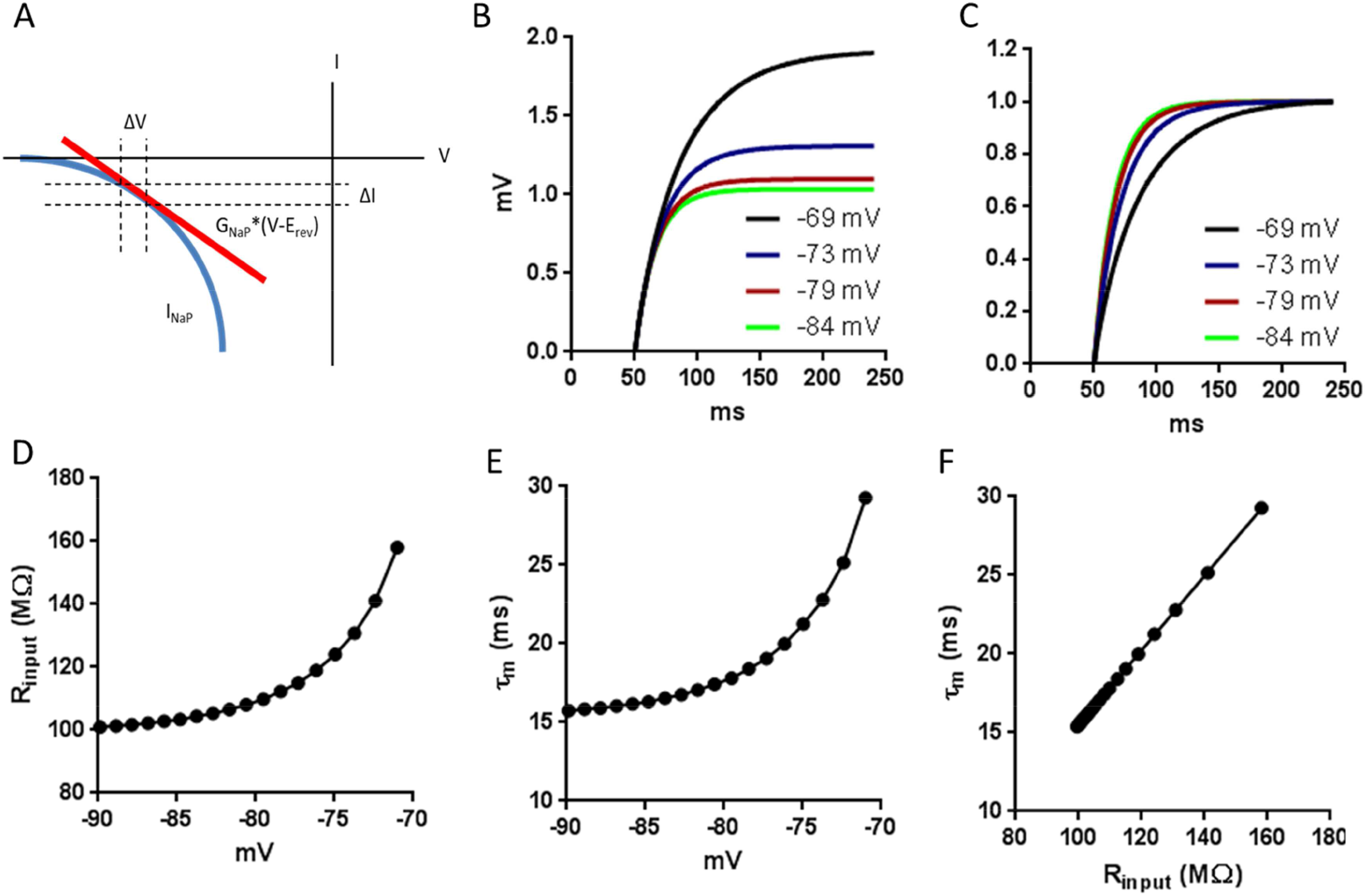
Computational simulations of voltage dependence of R_in_ and τ_m_. A. Approximation of instantaneous IV curve of I_NaP_ (blue) using a linear negative conductance (red) within the region limited by the dotted lines. B. Voltage dependence of membrane potential changes in response to a current of 10 pA C. Voltage dependence of the rise time of the responses in B. The graphs are normalized to the maximal response. D. Plot of the voltage dependence of R_in_ E. Plot of the voltage dependence of τ_m_. F. Correlation of τ_m_ vs R_in_. R^2^ = 1.

### I_NaP_ prolongs EPSP decay in a neuron model

In order to gain some insight about how I_nap_ shapes the EPSP and the role of the activation and deactivation kinetics of I_NaP_, we conducted computational simulations of I_NaP_ and EPSPs. Figures 2A shows examples of an EPSC produced by a triangular current (shown below) in the absence and in the presence of I_NaP_, with 2 activation time constants. Figure 2B shows that in the presence of I_nap_ with fast activation (τ_act_ = 0.1 ms), the EPSP amplitude is increased with membrane potential depolarization. In contrast, Figure 2C shows that in the presence of I_nap_ with slow activation (τ_act_ = 100 ms), the EPSP amplitude decreases with depolarization of the membrane potential. Figure 2D shows I_nap_ with its real fast activation and deactivation values (Carter et al., 2012) (τ = 0.1 ms), increases EPSP area in a voltage dependent manner. These both amplifies and slows down EPSPs, and that a fast activation is essential for its amplification and that even with a fast deactivation I_NaP_ can prolong the EPSP.

**Figure 2.**
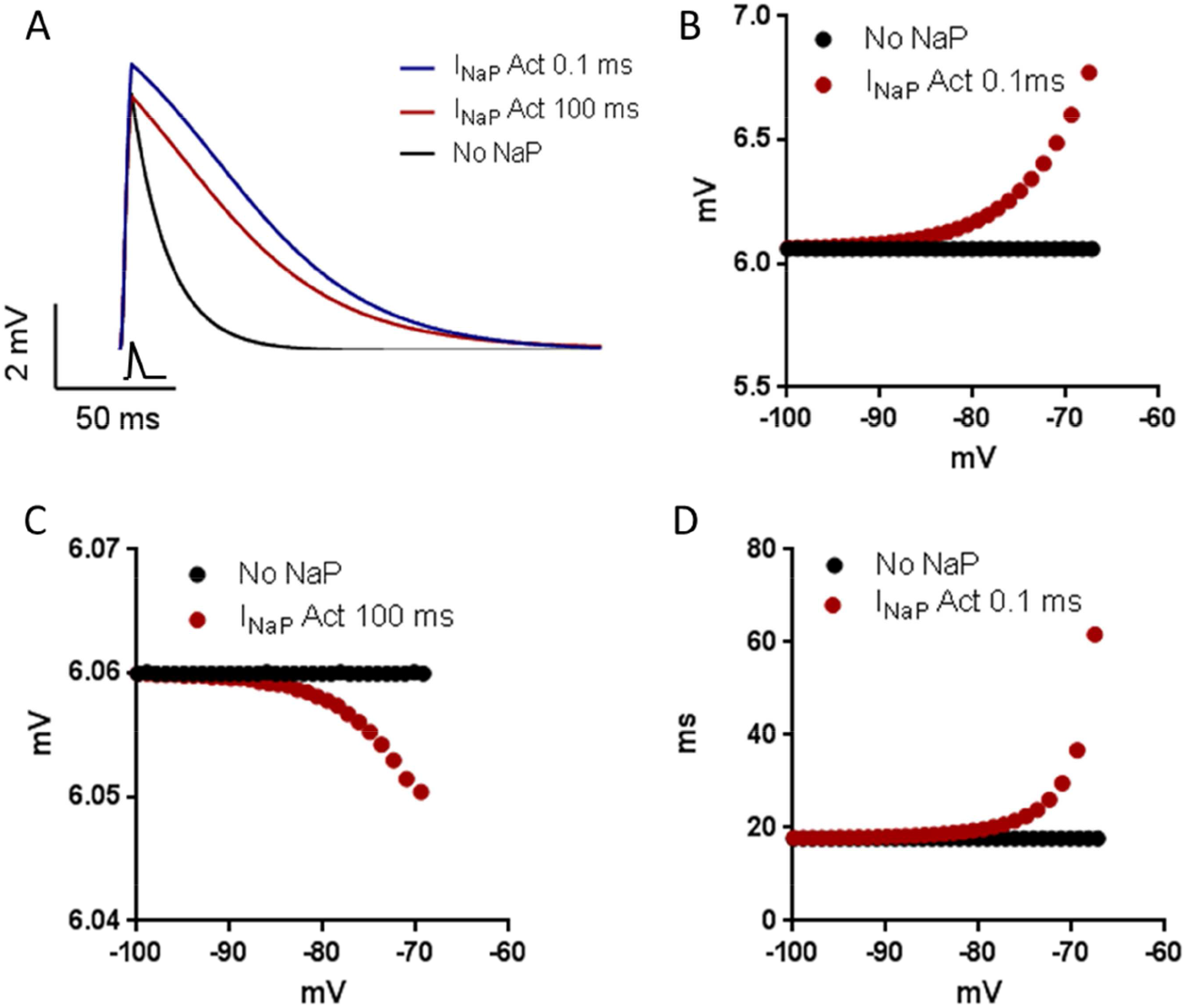
A. EPSP representative traces obtained from computational simulations for I_nap_ with fast activation (τ_act_ = 0.1 ms) and slow activation (τ_act_ = 100 ms). V = - 67 mV. aEPSC is drawn below the trace. B. Comparing EPSP amplitude for only linear current and with I_nap_ with fast activation (τ_act_ = 0.1 ms) for different membrane potentials. C. Comparing EPSP amplitude for only linear current and with I_nap_ with slow activation (τ_act_ = 100 ms) for different membrane potentials. D. EPSP area normalized by the amplitude with I_nap_ fast activation and deactivation (τ = 0.1 ms)

### G_NaP_ is composed mainly by its derivative component

Hippocampal CA1 pyramidal cells express I_NaP_ and this current produces a region of negative slope conductance (Yamada-Hanff & Bean, 2015; French et al., 1999; Carter et al., 2012) as can be seen in figure 3A. Application of a slow ramp (15 mV/s) produced an inward current which start to develop approximately at–85 mV and peaked around −40 mV. This current was abolished by TTX (100 nM) and the TTX-sensitive current had a peak of −537 ± 42 pA at the potential of −40 mV. Assuming a calculated sodium reversal potential E_Na_ = 70 mV, we calculated the NaP chord conductance (g_nap_) and it was 4.9 ± 0.4 nS at −40 mV. I_NaP_ had a half-activation potential of −55.6 ± 0.9 mV and a slope factor of 6.2 ± 0.4 mV (n = 10, Figure 3A), in line with previous reports (Yamada-Hanff & Bean, 2015, Carter et al., 2012).

We then measured G_nap_ by the first derivative of the current (dI/dV) and accordingly foud that it is is negative and voltage dependent, increasing with depolarization (p = 0.0008; One way repeated measures ANOVA; Figure 3B). On the other hand, the chord conductance g_nap_ is positive and equally voltage dependent, (p = 0.0045; One way repeated measures ANOVA; Figure 3C). But when we measured the derivative term G_nap_^Der^, using equation 9, we found that it is also negative and voltage dependent, increasing with depolarization (p = 0.0003; One way repeated measures ANOVA) (Figure 3D). Figure 3E summarizes the absolute values of G_nap_, g_nap_ and G_nap_^Der^ at different potentials showing that the major contribution of G_nap_ comes mostly from the G_nap_^Der^.

**Fig 3.**
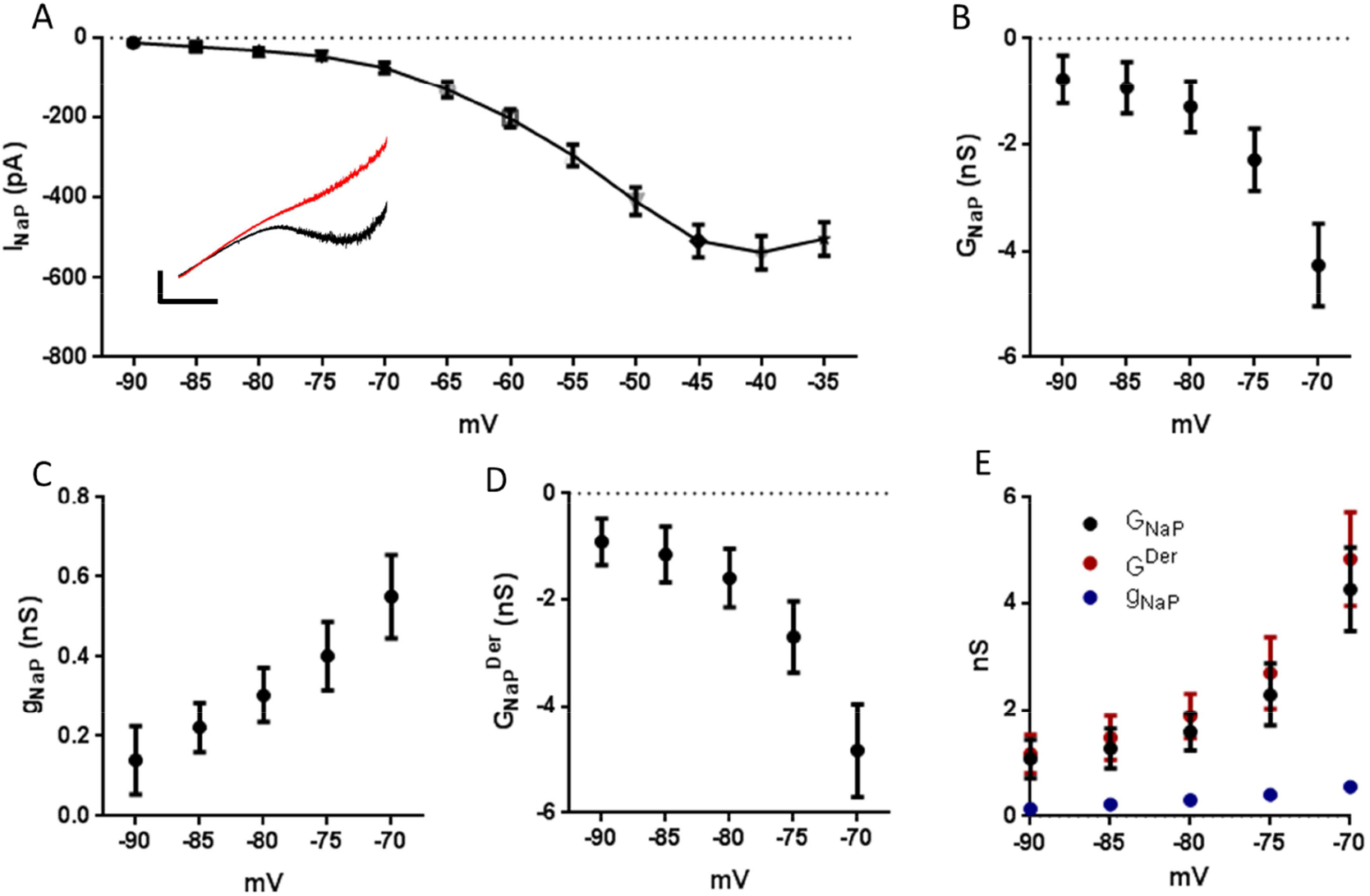
A. Average for I-V curves of persistent sodium current (I_NaP_) (n = 10) obtained by the subtraction of currents elicited by slow ramp command in a control (black) situation and after the perfusion with TTX (red). Inset. Representative traces. B. I_NaP_ slope conductance (G_nap_) determined from I-V curves in A (n = 9). C. I_NaP_ chord conductance (g_nap_) (n = 10). D. Derivative term of the slope conductance (G_nap_^Der^) obtained Der from B, C and D. subtracting G_nap_ from g_nap_ (n = 9). E. Summary of absolute values of G_nap_, g_nap_ and G_nap_

### The negative slope conductance of I_NaP_ is sufficient to increase the R_in_

The progressive activation of I_NaP_ during a ramp depolarization potentiates the depolarization of the membrane as can be seen in figure 4A, which is blocked by TTX (100 nM). In accordance to our theoretical prediction and to previous observations (Economo et al., 2014, Yamada-Hanff et al., 2015), R_in_ measured in current clamp (Figure 4B) is voltage dependent before and after TTX application. However, TTX decreases significantly the voltage-dependence of R_in_ (p<0.0001 Two way repeated measures ANOVA n = 6), by reducing the voltage effect produced by the more depolarized potentials (−75 and −70 mV, p<0.05 Tukey’s multiple comparison test). Since the mean resting membrane potential (RMP) was −78.3 ± 1.1 mV, these results show that the effect of I_NaP_ on R_in_ is strong only for potentials above the RMP. Similar results were obtained in voltage-clamp (not shown).

To demonstrate that TTX decreases the R_in_ by inhibiting of the I_NaP_ negative slope conductance we conducted experiments with TTX and applied to the neuron an artificial I_NaP_ (aI_NaP_) using dynamic-clamp. Figure 4B shows that aI_NaP_ restores the voltage dependency of R_in_ after TTX (Two way repeated measures ANOVA, n = 6), by increasing the effect of more positive potentials (−75 mV Tukey’s multiple comparison test).

To confirm that the ability for I_NaP_ to increase R_in_ is related to its negative slope conductance, we injected artificial linear currents in pyramidal cells using dynamic clamp. We then tested two linear currents: One with a positive conductance and the other with a negative conductance. Where the current with positive conductance represents g_NaP_ (ag_NaP_), meanwhile the current with negative conductance represents G_NaP_ (aG_NaP_) (Figure 4E).

Figure 4C shows that aG_NaP_ was able to restore the high values of R_in_ associated with the presence of I_NaP_ (ttx vs aG_NaP_, Two way repeated measures ANOVA, n = 9). On the other hand, ag_NaP_ did not change the R_in_ (Figure 4D). In Figure 4E (right), we compare the current values in the range between −80 and −70 mV for the aG_NaP_, ag_NaP_ and aI_NaP_. Despite that, within this range, the current of the ag_NaP_ was higher than the current of the aG_NaP_ and the aI_NaP_, only the latter currents were effective influencing R_in_, showing that R_in_ reflects the rate of the change of the currents (i.e. the slope conductance) and not its absolute values (i.e. instantaneous currents).

**Fig 4.**
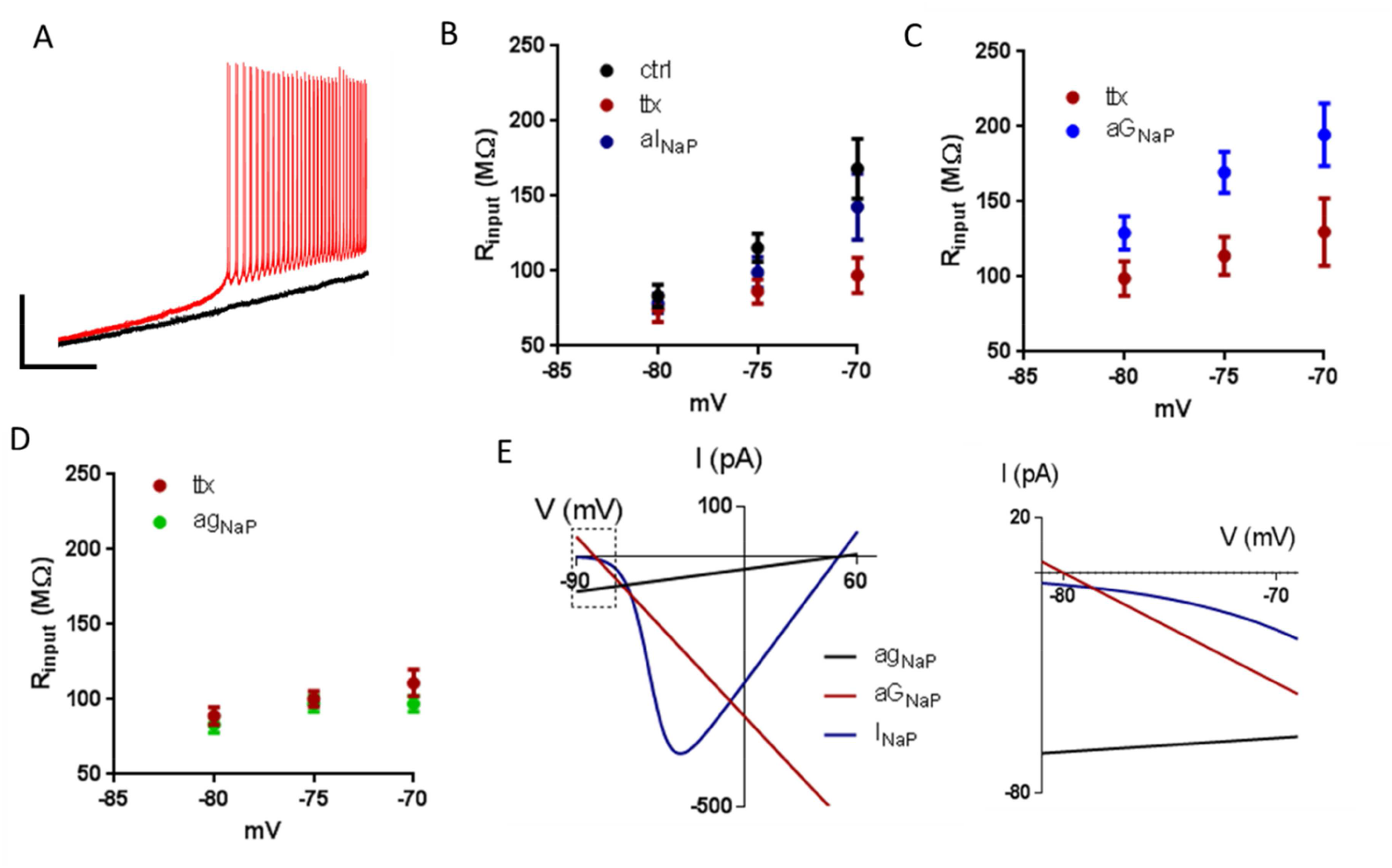
A. Ramp recorded in the current-clamp mode for the control case (red) and after the application of TTX (black). B. Voltage dependence of R_in_ before and after the application of TTX and with the injection of aI_NaP_ (n = 6). C. Voltage dependence of R_in_ after the application of TTX and with the injection of aG_NaP_ (n = 9). D. Voltage dependence of R_in_ after the perfusion with TTX and with the injection of ag_NaP_ (n = 8). E. IV curves of ag_NaP_, aG_NaP_ and aI_NaP_ injected using dynamic-clamp (Left). Expanded plot focusing on the membrane potential region which the aG_NaP_ was injected (V_m_ > −80 and V_m_ < −70 mV) (right). It is noted that all the three currents are depolarizing.

### Membrane time constant is voltage dependence and is determined by the voltage-dependence of R_in_

In accordance to our theoretical prediction and to what was observed by others (Yamada-Hanff et al., 2015), we observed that τ_m_ is voltage dependent and increases with depolarization, an effect abolished by TTX application (Figure 5A and Figure 5B, Two way repeated measures ANOVA, n = 11). Similarly to what observed with R_in_, TTX application decreases significantly the τ_m_ in more depolarized potentials (−75 mV, p<0.05 Sidak’s multiple comparison test). We then tested if applying aG_NaP_ could increase τ_m_ in the presence of TTX. Figure 5C shows that aG_NaP_ increases τ_m_ independently of voltage to values similar to τ_m_ measured without TTX at −75 mV (Two way repeated measures ANOVA, n = 10; Figure 5C). Accordingly to an effect dependent on the negative slope conductance, ag_NaP_ did not change the τ_m_ (Figure 5D, Two way repeated measures ANOVA, n = 11). Similarly, as already discussed above with R_in_, aG_NaP_ and not ag_NaP_ was effective influencing τ_m_, showing that τ_m_ reflects the rate of the change of the currents (i.e. the slope conductance) and not its absolute values (i.e. instantaneous currents).

Accordingly to the predicted by the cable theory, the mathematical relation τ_m_ = C* R_in_ must be valid for a passive membrane. However we do not know if for our situation of a membrane containing leak and NaP channels this is still valid. So we correlated the values of R_in_ and τ_m_ for all the membrane potentials for the control group and observed a positive correlation between R_in_ and τ_m_ (p < 0.0001, R^2^ = 0.556 n= 12 neurons). The equation slope determined from the linear regression was 196 pF what is in line with the capacitance values determined by others in CA1 pyramidal neurons (Tamagnini et al., 2015).

**Fig 5.**
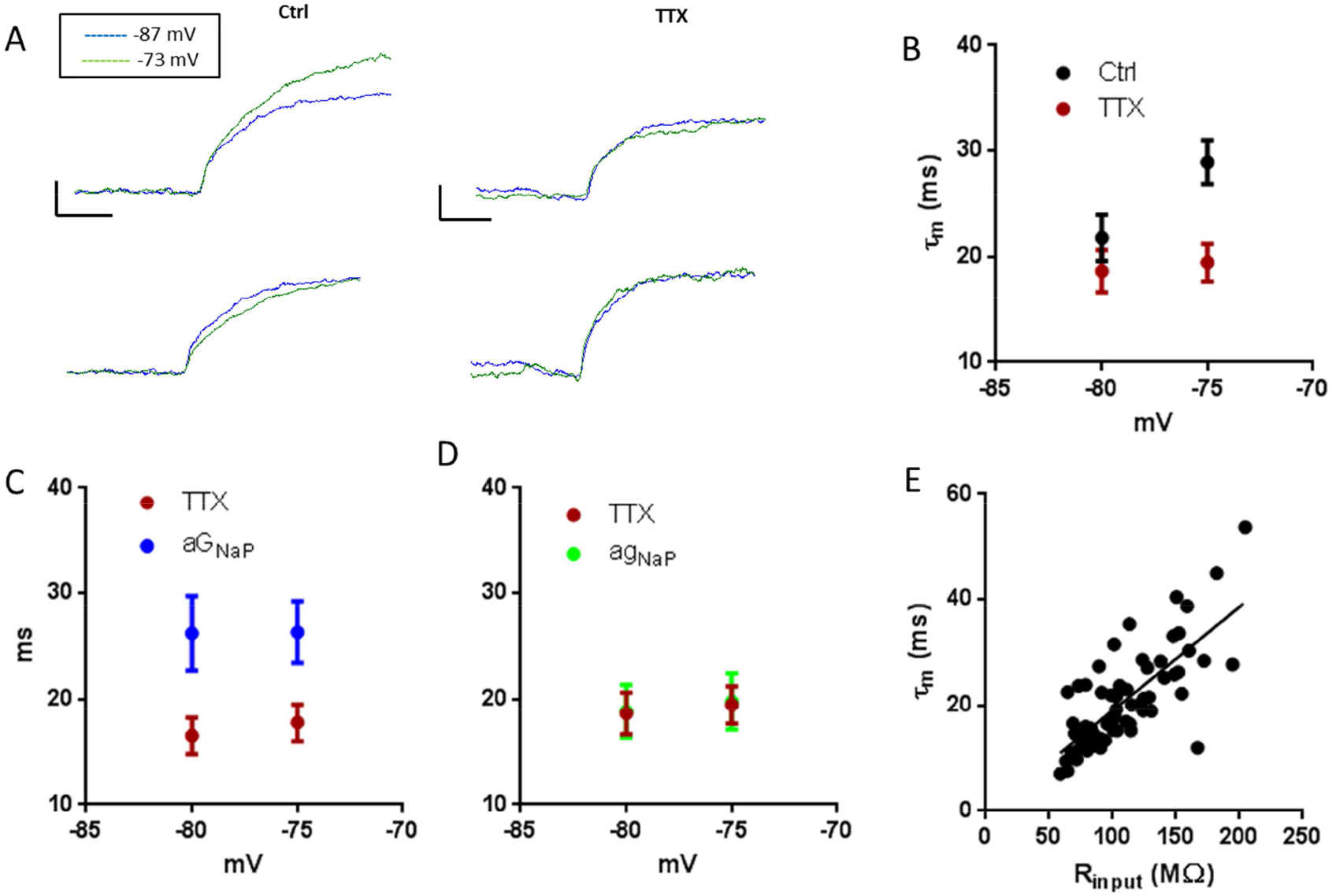
A. Representatives diagrams of the same neuron, control and TTX. Voltage change increases with depolarization in the control group but not in TTX group (top). τ_m_ increases with depolarization in the control group but not in TTX group (bottom, same traces from top but normalized). The current injected was 20 pA. Bar sizes: vertical: 1 mV, horizontal: 20 ms. B. Voltage dependence of τ_m_ for control and TTX group (n = 11). C. Voltage dependence of τ_m_ after TTX application and with the injection of aG_NaP_. (n = 10). D. Voltage dependence of τ_m_ after TTX application and with the injection of ag_NaP_. (n = 11). F. Linear regression of R_in_ vs τ_m_ for control group for all the membrane potentials.

### The negative slope conductance explains amplification of EPSPs

We then tested if I_nap_ could amplify an artificial EPSP (aEPSP) amplitude and decay time, measuring the amplitude and area of aEPSPs for different membrane potentials before and after TTX application. The aEPSP amplitude is voltage-dependent and increased slightly with depolarization by 5% when comparing −85 with −70 mV. Application of TTX abolished this voltage-dependency. Figure 6A and Figure 6B show that the aEPSP area is also voltage-dependent and increased considerably with depolarization by 30% when comparing −85 with −70 mV. Application of TTX abolished this voltage-dependency confirming the role of I_NaP_ (Two way repeated measures ANOVA, n = 10).

We then wanted to know if the prolongation of the aEPSP by I_NaP_ was caused by its negative slope conductance G_NaP_. For this we applied an artificial G_NaP_ in the presence of TTX in order to know if can amplify and prolong the aEPSPs. The injection of aG_NaP_ increases the EPSP amplitude in a voltage independent manner in accordance to what observed with R_in_ previously. To quantify the effect of aG_NaP_ on the decay of the aEPSP, we calculated the area of the normalized aEPSPs. As can be seen in Figure 6C and Figure 6D, the injection of aG_NaP_ increases the EPSP normalized area by the amplitude in a voltage independent manner (Two way repeated measures ANOVA, n =9). These results strongly suggest that I_nap_ EPSP amplification is mainly due to its negative slope conductance.

**Fig 6.**
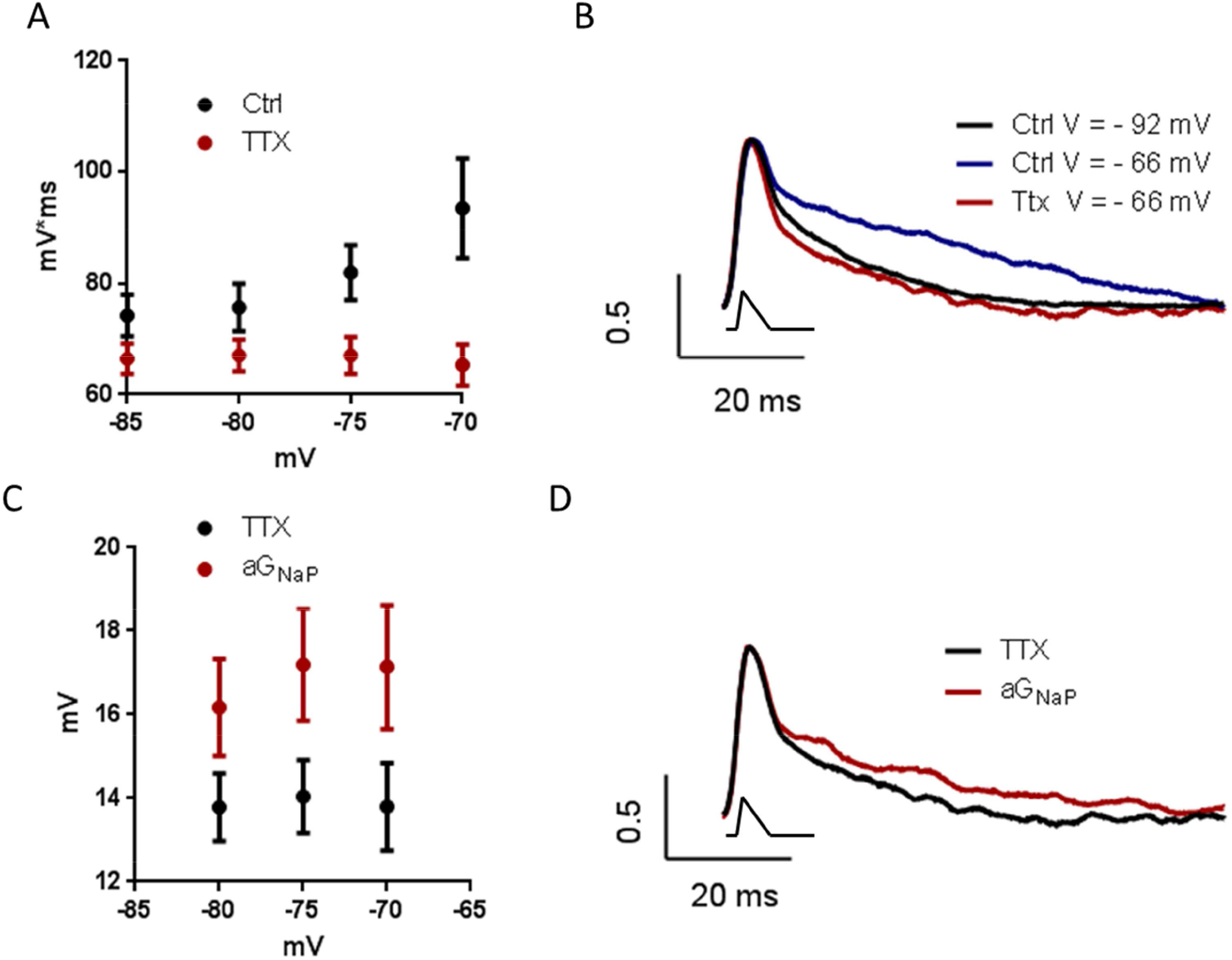
A. EPSPs area comparing the control vs TTX. B. EPSP representative traces. EPSP normalized by the amplitude for control for depolarized (V = −66 mV) and hyperpolarized membrane potentials (V = −92 mV) and for TTX for depolarized membrane potential (V = −66 mV). aEPSC is drawn below the traces. C. EPSPs normalized area by the amplitude comparing TTX vs the injection of aG_NaP_. D. EPSP representative traces. EPSP normalized by the amplitude for TTX vs the injection of aG_NaP_ for a depolarized membrane potential (V = −66 mV). aEPSC is drawn below the traces.

## DISCUSSION

In this work we developed a new analytical solution of the slope conductance of the non-inactivating voltage dependent currents in order to understand the mechanisms by which I_NaP_ activation increases R_in_ and prolongs τ_m_. Using mathematical, computational and experimental approaches, we established the biophysical mechanism by which a negative slope conductance increases R_in_ and τ_m_. Our results show that the amplification of the amplitude and the prolongation of the decay time of subthreshold EPSPs by I_NaP_ are basically caused by its negative slope conductance, and it is not related to its deactivation kinetics or other voltage-dependent conductance activated by I_NaP_. Additionally we showed that I_NaP_ is well suited for this because of its very fast activation kinetics, which produces an almost instantaneous current, which behaves during its activation phase like a linear negative conductance.

Accordingly to previous observations we observed that under presence of the endogenous I_NaP_, R_in_ and τ_m_ increases in a voltage dependent manner, an effect abolished by TTX application (Wilson 2005, Haj-Dahmane et al., 1996, Yamada-Hanff et al., 2013, Fernandez et al., 2015, Economo et al., 2014, Yamada-Hanff et al., 2015, Jacobson et al., 2005, Yaron-Jakoubovitch et al., 2008, Boehlen 2012, Waters et al., 2006, Buchanan et al., 1992). These previous reports suggested that this paradoxical increase in R_in_ produced by the activation of NaP current were caused by a negative slope conductance region generated by I_NaP._. On the other hand, they were not conclusive about the effect on τ_m_. Our dynamic clamp experiments showed that injection of a pure negative linear conductance (aG_NaP_) increased the R_in_ and τ_m_, validating the hypothesis that the ability for I_NaP_ to increase R_in_ and τ_m_ is related to its negative slope conductance. Thus the electrical behavior of I_NaP_ is sufficient to explain the observed increase of R_in_ and τ_m_, and not a result of other conductance activated by Na^+^ or other subthreshold conductance.

Using mathematical and computational simulations we determined that because I_NaP_ has a fast activation/deactivation kinetics, when compared with the cell membrane kinetics, I_NaP_ reaches is steady state so fast that I_NaP_ is be able to control the τ_m_ via its steady state slope conductance G_NaP_. Thus, we believe that the contribution of I_NaP_ to the τ_m_ is via its negative slope conductance G_NaP_. Our computational simulations showed that τ_m_ is directly proportional to R_in_, independently the membrane potential, suggesting that the relation τ_m_ = C_m_*R_in_ is valid. This is in agreement to the behavior of passive membranes in accordance with the cable theory, even being I_NaP_ a voltage-dependent current.

Previous works established that the slope conductance of non-inactivating voltage dependent currents has two terms: a passive and a derivative (Koch 1998, Wessel et al., 1999, Ferreira et al., 1985). However, it was unknown how much is the contribution of each component to the slope conductance of I_NaP_. We found that the derivative term of I_NaP_ 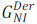 is always negative while the passive term I^NaP^ 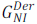 is always positive. We also found that the magnitude of 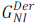 was much bigger than 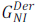, resulting in a negative G_nap_ mainly dominated by the contribution of its derivative term. In fact, 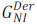 was at least 10 times bigger than 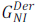, so more than 90 % of the G_NaP_ comes from 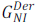 magnitude. This is because the driving force is a multiplicative factor of 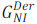 but not of 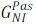, and because the subthreshold membrane potential are distant from the sodium reversal potential (+70 mV), creating a large driving force for sodium (120 mV).

The paradox of blocking I_NaP_ decreases R_in_ and τ_m_ is a consequence of the assuming a direct correlation of conductance with permeability. The classical interpretation of the membrane input resistance is based on the assumption that it is inversely related to the conductance (permeability) of the channel. So that the most logical interpretation when the input resistance increases or decreases is that the channel is closed or opened. The definition of the slope conductance is not directly connected with the description of membrane conductance being defined as an increase in the membrane ionic permeability. In order to explain the input resistance and its voltage dependent behavior, the slope conductance is more appropriate than the chord conductance, which has a direct correspondence to the membrane ionic permeability. The I_NaP_ I-V curve provides a good example of the difference between the slope and chord conductances. The I_NaP_ slope conductance is negative at subthreshold membrane potentials, but its chord conductance is always positive at every value of membrane potential. The ohmic conductance of the channel can only be obtained by computing the chord conductance at each voltage. The chord conductance, therefore, is most appropriate for characterizing the ionic permeability of the membrane while the slope conductance reflects an electric property directly affecting the membrane input resistance. Earlier attempts have been made in this direction. Izhikevich (2007) and Hutcheon et al., (2000) classified the voltage dependent currents in two categories: amplifying and resonant currents. Amplifying currents (e.g. I_NaP_) exhibit negative slope conductance meanwhile resonant currents (e.g. I_h_) only exhibit positive slope conductance, affecting the membrane input resistance differentially.

Here, we propose that for I_NaP_, it is the slope conductance and not the chord conductance which is more relevant to produce its influence on R_in_ and τ_m_. In such paradigm, slope conductances (positive and negative) from each channel (linear and voltage-dependent) is simply added algebraically, and it is this sum that determine R_in_ and τ_m_, following a simple mathematical rule: R_in_ = 1/ΣG and τ_m_ = R_in_*C, (where C is the membrane capacitance). It is important to highlight that this paradigm is only valid for voltage dependent currents with fast kinetics. From the point of view of this, the anomalous increase of R_in_ by the activation of the NaP channels is no longer a counterintuitive result, and it is easily explained by an electric circuit as a consequence of positive and negative slope conductances in parallel which can be summed algebraically (Eeckman 2012). Thus, in the near threshold region, where I_h_ contributes less than the leak current, G_NaP_ opposes the positive slope conductance of the leak current (Yamada-Hanff et al., 2015), which leads to a decrease of the total positive slope conductance, leading to an increase of the R_in_ of the neuron.

These concepts can be extended to understand the effect of other ionic currents, besides I_NaP_, that also present a negative slope conductance regions, such as some Ca^++^ voltage dependent channels, NMDA receptors and inward rectifier potassium currents. For instance, the anomalous increase of the R_in_ in neurons that is observed also during the activation of NMDA receptors, can be explained as a consequence of a negative slope conductance region (Crunelli et al., 1984, Moore et al., 1995, Sutor et al., 1987, MacDonald et al., 1982, Koch 1998). In addition, it has been observed a prolongation of the EPSP decay time due to Ca channels opening (Prescott& De Koninck, 2005).

As we observed in our results, the effect of I_NaP_ on the excitability of the neuron membrane is beyond its effects on the R_in_ and τ_m_. For instance, a well-recognized phenomenon is the amplification of the EPSPs and IPSPs by the I_NaP_ (Stuart et al., 1995, Schwindt et al., 1995, Stuart 1999, Fricker et al., 2000, Hardie et al, 2006). It is well known that I_NaP_ amplifies the PSPs by increasing their amplitude and prolonging their decay. We observed this same amplification in our experiments. In addition, we observed the same amplification when injected a linear negative conductance after TTX application. These results support the hypotheses that I_nap_ amplification of EPSP is mainly due to its negative slope conductance. Since artificial EPSC were injected through the recording pipette placed in the neuron soma, our results suggest that the effects observed in this work on the EPSP amplification are mainly due to the recruitment of sodium channels placed in the soma (Andreasen & Lambert, 1999). Also, our results are in agreement with a recent developed theory called the quasi-active cable approximation that showed that amplifying currents such as I_NaP_ amplifies the EPSP amplitude and prolong its decay time (Remme & Rinzel 2011).

Thus, our results contribute to the understanding of the mechanisms behind the amplification of PSPs, since increasing of the PSP amplitude and prolonging of their decay can be interpreted as a straightforward result of the increase of the R_in_ and τ_m_ by I_NaP_. For instance, in a previous study, Rotaru et al., 2015 reported that cells’ input resistance and the cells’ membrane time constant shape partially PSP amplitude and time course. More evidence supporting our hypothesis that it is the negative slope conductance of I_NaP_ that prolongs the EPSP decay is found in Zsiros, V., & Hestrin 2005, that showed that injecting a positive conductance but not a current shortens the prolongation of the EPSP decay enhanced by I_NaP_. Furthermore, in an extreme case, Farries et al., 2010 showed that I_NaP_ negative slope conductance can oppose the positive slope conductance due to the other subthreshold currents, thus creating a wide region with zero slope conductance that establishes an infinite time constant and avoid the decay phase of EPSPs.

Computational simulations showed that I_NaP_ fast activation is necessary to amplify the EPSP amplitude, meanwhile slow activation leads to decrease of the EPSP amplitude. These results can be explained since when the activation is fast, G_NaP_ influences the dynamics and amplifies the EPSP, meanwhile when the activation is slow, g_NaP_ but not G_NaP_ influences the dynamics and diminishes the EPSP (Remme & Rinzel, 2011). Strikingly, computational simulations also showed that a fast deactivation kinetics (τ_deact_ = 0.1 ms) prolongs the EPSP decay time when compared with no I_nap_. These results showed that, although counterintuitive, fast I_nap_ deactivation kinetics slows down EPSP decay time. These results are in line with experimental data showing that I_nap_ kinetics in CA1 pyramidal cells is extremely fast (Carter et al., 2012). These results support our claim that it is the I_NaP_ negative slope conductance and not an slow channel kinetics the cause of the prolongation of the EPSP decay time.

It is believed that activation of inward currents amplify EPSPs by increasing their amplitude and prolonging their decay, while the activation of outward currents may counter the effects of inward currents on EPSP shape (Fricker et al., 2000). However, that view in which the classification of the current as inward or outward determines their effects on the PSPs shape has flaws for several reasons. For instance, the activation of the h current, that is an inward current, decreases EPSPs and IPSPs amplitude and reduces their decay (Magee 1998, Hardie et al, 2006). Moreover, I_NaP_, that is also an inward current, amplifies IPSPs (Stuart 1999, Hardie et al, 2006), but none known outward current amplifies IPSPs. Also, a leak current could be inward or outward according to its reversal potential, however leak currents always decrease PSPs amplitude and reduces time decay. Thus, in order to reconcile the experimental data and the theoretical knowledge, we emphasized that a more appropriate view requires the dissociation of two main concepts: 1) the direction of the current flux of the channel, i.e., inward or outward currents and 2) the slope conductance of the channel. Following that line of thought, a current with a negative slope conductance and fast kinetics, it increases the input resistance and amplifies both EPSPs and IPSPs regardless it is an inward or outward current. The opposite happens with a current with a positive slope conductance.

In summary, we propose that the main role of the I_NaP_ in the membrane excitability is the increase of R_in_ due to its negative slope conductance. The other phenomena are a consequence of that increase of R_in_, such as: increase of τ_m_, EPSP amplitude amplification and decay time prolongation.

## REFERENCES

Aman TK, Grieco-Calub TM, Chen C, Rusconi R, Slat EA, Isom LL, Raman IM. (2009) Regulation of persistent Na current by interactions between beta subunits of voltage-gated Na channels. J Neurosci 29:2027–42.

Andreasen, M., & Lambert, J. D. (1999). Somatic amplification of distally generated subthreshold EPSPs in rat hippocampal pyramidal neurones. The Journal of Physiology, 519(1), 85–100.

Boehlen, A., Henneberger, C., Heinemann, U., & Erchova, I. (2012). Contribution of near-threshold currents to intrinsic oscillatory activity in rat medial entorhinal cortex layer II stellate cells. J Neurophysiol, 109:445–463.

Bose, A., Golowasch, J., Guan, Y., & Nadim, F. (2014). The role of linear and voltage-dependent ionic currents in the generation of slow wave oscillations. J Comput Neurosci, 37:229–242.

Buchanan, J. T., Moore, L. E., Hill, R., Wallén, P., & Grillner, S. (1992). Synaptic potentials and transfer functions of lamprey spinal neurons. Biol Cybernet, 67:123–131.

Carter, B. C., Giessel, A. J., Sabatini, B. L., & Bean, B. P. (2012). Transient sodium current at subthreshold voltages: activation by EPSP waveforms. Neuron, 75:1081–1093.

Crill WE (1996) Persistent sodium current in mammalian central neurons. Annu Rev Physiol, 58:349–362.

Crunelli, V. & Mayer, M. L. (1984). Mg 2+ dependence of membrane resistance increases evoked by NMDA in hippocampal neurones. Brain Res, 311:392–396.

Dagostin, A. A., Lovell, P. V., Hilscher, M. M., Mello, C. V., & Leão, R. M. (2015). Control of phasic firing by a background leak current in avian forebrain auditory neurons. Frontiers in cellular neuroscience, 9.

Dallérac, G., Chever, O., & Rouach, N. (2014). How do astrocytes shape synaptic transmission? Insights from electrophysiology. Front Cell Neurosci, 7:159.

Deisz, R. A., Fortin, G., & Zieglgänsberger, W. (1991). Voltage dependence of excitatory postsynaptic potentials of rat neocortical neurons. J Neurophysiol, 65:371–382.

Del Negro, C.A., Koshiya, N., Butera, R.J. & Smith, J.C. (2002) Persistent sodium current, membrane properties and bursting behavior of pre-Boltzinger complex inspiratory neurons in vitro. J. Neurophysiol., 88, 2242–2250

Du, Y., Kiyoshi, C. M., Wang, Q., Wang, W., Ma, B., Alford, C. C., Zhong, S., Wan, Q., Chen, H., Lloyd, E. E., Bryan Jr., R. M. & Zhou, M.. (2016). Genetic deletion of TREK-1 or TWIK-1/TREK-1 potassium channels does not alter the basic electrophysiological properties of mature hippocampal astrocytes in situ. Front Cell Neurosci, 10:13.

Economo, M. N., Martínez, J. J., & White, J. A. (2014). Membrane potential-dependent integration of synaptic inputs in entorhinal stellate neurons. Hippocampus, 24:1493–1505.

Enyedi, P., & Czirják, G. (2010). Molecular background of leak K+ currents: two-pore domain potassium channels. Physiol Rev, 90:559–605.

Farries, M. A., Kita, H., & Wilson, C. J. (2010). Dynamic spike threshold and zero membrane slope conductance shape the response of subthalamic neurons to cortical input. The Journal of Neuroscience, 30(39), 13180–13191.

Fernandez, F. R., Malerba, P., & White, J. A. (2015). Non-linear membrane properties in entorhinal cortical stellate cells reduce modulation of input-output responses by voltage fluctuations. PLoS Comput Biol, 11:e1004188.

Ferreira, H. G., & Marshall, M. W. (1985). The Biophysical Basis of Excitability. Cambridge University Press.

French, C. R., Sah, P., Buckett, K. J., & Gage, P. W. (1990). A voltage-dependent persistent sodium current in mammalian hippocampal neurons. J Gen Physiol, 95:, 1139–1157.

Fricker, D., & Miles, R. (2000). EPSP amplification and the precision of spike timing in hippocampal neurons. Neuron, 28:559–569.

Ghigliazza, R. M., & Holmes, P. (2004). Minimal models of bursting neurons: How multiple currents, conductances, and timescales affect bifurcation diagrams. SIAM J App Dyn Syst, 3:636–670.

Haj-Dahmane, S., & Andrade, R. (1996). Muscarinic activation of a voltage-dependent cation nonselective current in rat association cortex. J Neurosci, 16:3848–3861.

Hardie, J. B., & Pearce, R. A. (2006). Active and passive membrane properties and intrinsic kinetics shape synaptic inhibition in hippocampal CA1 pyramidal neurons. J Neurosci, 26:8559–8569.

Hutcheon, B., & Yarom, Y. (2000). Resonance, oscillation and the intrinsic frequency preferences of neurons. TINS, 23:216–222.

Izhikevich, E. M. (2007). Dynamical Systems in Neuroscience. MIT Press.

Jacobson, G. A., Diba, K., Yaron-Jakoubovitch, A., Oz, Y., Koch, C., Segev, I., & Yarom, Y. (2005). Subthreshold voltage noise of rat neocortical pyramidal neurones. J Physiol, 564:145–160.

Klink, R., & Alonso, A. (1997). Ionic mechanisms of muscarinic depolarization in entorhinal cortex layer II neurons. J Neurophysiol, 77:1829–1843.

Koch, C. (1998). Biophysics of Computation: Information Processing in Single Neurons. Oxford University Press.

König, P., Engel, A. K., & Singer, W. (1996). Integrator or coincidence detector? The role of the cortical neuron revisited. TINS, 19:130–137.

Leão RM, Kushmerick C, Pinaud R, Renden R, Li GL, Taschenberger H, Spirou G, Levinson SR, von Gersdorff H. (2005) Presynaptic Na^+^ channels: locus, development, and recovery from inactivation at a high-fidelity synapse. J Neurosci. 25:3724–38.

Leao, R. M., Li, S., Doiron, B., & Tzounopoulos, T. (2012). Diverse levels of an inwardly rectifying potassium conductance generate heterogeneous neuronal behavior in a population of dorsal cochlear nucleus pyramidal neurons. Journal of neurophysiology, 107(11), 3008–3019.

MacDonald, J. F., Porietis, A. V., & Wojtowicz, J. M. (1982). L-Aspartic acid induces a region of negative slope conductance in the current-voltage relationship of cultured spinal cord neurons. Brain Res, 237:248–253.

Magee, J. C. (1998). Dendritic hyperpolarization-activated currents modify the integrative properties of hippocampal CA1 pyramidal neurons. J Neurosci, 18:7613–7624.

Magistretti J, Ragsdale DS, Alonso A (1999) High conductance sustained single-channel activity responsible for the low-threshold persistent Na^+^ current in entorhinal cortex neurons. J Neurosci 19:7334–7341.

Moore, L. E., Buchanan, J. T., & Murphey, C. R. (1995). Localization and interaction of N-methyl-D-aspartate and non-N-methyl-D-aspartate receptors of lamprey spinal neurons. Biophys J, 68:96–103.

Moore, L. E., Buchanan, J. R. & Murphey, C. R. (2012). Anomalous increase in membrane impedance of neurons during NMDA activation. In: Eeckman, F. (Ed.). (2012). Computation in Neurons and Neural Systems. Springer Science & Business Media. pp. 21–26.

Nisenbaum, E.S. and Wilson, C.J. (1995) Potassium current responsible for inward and outward rectification in rat neostriatal spiny projection neurons. J Neurosci 15:4449–4463.

Pennartz, C. M. A., Bierlaagh, M. A., & Geurtsen, A. M. S. (1997). Cellular mechanisms underlying spontaneous firing in rat suprachiasmatic nucleus: involvement of a slowly inactivating component of sodium current. Journal of Neurophysiology, 78(4), 1811–1825.

Prescott, S. A., & De Koninck, Y. (2005). Integration time in a subset of spinal lamina I neurons is lengthened by sodium and calcium currents acting synergistically to prolong subthreshold depolarization. The Journal of neuroscience, 25(19), 4743–4754.

Raman, I. M., Gustafson, A. E., & Padgett, D. (2000). Ionic currents and spontaneous firing in neurons isolated from the cerebellar nuclei. The Journal of Neuroscience, 20(24), 9004–9016.

Qu Y, Curtis R, Lawson D, Gilbride K, Ge P, DiStefano PS, Silos-Santiago I, Catterall WA, Scheuer T (2001) Differential modulation of sodium channel gating and persistent sodium currents by the β1, β2, and β3 subunits. Mol Cell Neurosci 18:570–580.

Remme, M. W., & Rinzel, J. (2011). Role of active dendritic conductances in subthreshold input integration. Journal of computational neuroscience, 31(1), 13–30.

Rotaru, D. C., Olezene, C., Miyamae, T., Povysheva, N. V., Zaitsev, A. V., Lewis, D. A., & Gonzalez-Burgos, G. (2015). Functional properties of GABA synaptic inputs onto GABA neurons in monkey prefrontal cortex. Journal of neurophysiology, 113(6), 1850–1861.

Schwindt, P. C., & Crill, W. E. (1995). Amplification of synaptic current by persistent sodium conductance in apical dendrite of neocortical neurons. J Neurophysiol, 74:2220–2224.

Sutor, B., Jordan, W., & Zieglgänsberger, W. (1987). Evidence for a magnesium-insensitive membrane resistance increase during NMDA-induced depolarizations in rat neocortical neurons in vitro. Neuroscience Let, 75:317–322.

Stuart, G., & Sakmann, B. (1995). Amplification of EPSPs by axosomatic sodium channels in neocortical pyramidal neurons. Neuron, 15:1065–1076.

Stuart, G. (1999). Voltage–activated sodium channels amplify inhibition in neocortical pyramidal neurons. Nat Neurosci, 2:144–150.

Stafstrom, C. E., Schwindt, P. C., & Crill, W. E. (1982). Negative slope conductance due to a persistent subthreshold sodium current in cat neocortical neurons in vitro. Brain Res, 236:221–226.

Stafstrom, C.E., Schwindt, P.C., Chubb, M.C., and Crill, W.E. (1985). Properties of persistent sodium conductance and calcium conductance of layer V neurons from cat sensorimotor cortex in vitro. J Neurophysiol, 53:153–170.

Stafstrom, C. E. (2007). Persistent sodium current and its role in epilepsy. Epilepsy Currents, 7(1), 15–22.

Surges, R., Freiman, T. M., & Feuerstein, T. J. (2004). Input resistance is voltage dependent due to activation of Ih channels in rat CA1 pyramidal cells. J Neurosci Res, 76: 475–480.

Tamagnini, F., Novelia, J., Kerrigan, T. L., Brown, J. T., Tsaneva-Atanasova, K., & Randall, A. D. (2015). Altered intrinsic excitability of hippocampal CA1 pyramidal neurons in aged PDAPP mice. Frontiers in cellular neuroscience, 9.

Thomson, A. M., Girdlestone, D., & West, D. C. (1988). Voltage-dependent currents prolong single-axon postsynaptic potentials in layer III pyramidal neurons in rat neocortical slices. J Neurophysiol, 60:1896–1907.

Waters, J., & Helmchen, F. (2006). Background synaptic activity is sparse in neocortex. J Neurosci, 26:8267–8277.

Wessel, R., Kristan, W. B., & Kleinfeld, D. (1999). Supralinear summation of synaptic inputs by an invertebrate neuron: dendritic gain is mediated by an “inward rectifier” K+ current. J Neurosci, 19:5875–5888.

Wilson, C. J. (2005). The mechanism of intrinsic amplification of hyperpolarizations and spontaneous bursting in striatal cholinergic interneurons. Neuron, 45:575–585.

Yamada-Hanff, J., & Bean, B. P. (2013). Persistent sodium current drives conditional pacemaking in CA1 pyramidal neurons under muscarinic stimulation. J Neurosci, 33:15011–15021.

Yamada-Hanff, J., & Bean, B. P. (2015). Activation of Ih and TTX-sensitive sodium current at subthreshold voltages during CA1 pyramidal neuron firing. J Neurophysiol, 114:2376–2389.

Yaron-Jakoubovitch, A., Jacobson, G. A., Koch, C., Segev, I., & Yarom, Y. (2008). A paradoxical isopotentiality: a spatially uniform noise spectrum in neocortical pyramidal cells. Front Cell Neurosci, 2:3.

Zsiros, V., & Hestrin, S. (2005). Background synaptic conductance and precision of EPSP-spike coupling at pyramidal cells. Journal of neurophysiology, 93(6), 3248–3256.

